# Azithromycin Derivatives to Mitigate Off-Target Inhibition of Autophagy and Retain Beneficial Host Directed Effects

**DOI:** 10.64898/2026.09.01.748460

**Authors:** Duc Q Doan, Shawn WH Liu, Samuel Clark, Sylvia A Sapula, Thomas Crowhurst, Paul N Reynolds, Benjamin T Kopp, Anton Blencowe, Eugene Roscioli

**Author notes:** Authors contributed equally (Anton Blencowe, Eugene Roscioli). Correspondence: Eugene Roscioli, Adelaide Health and Medical Science Building Corner of North Terrace and George St Adelaide, 5005, South Australia Australia.

## Abstract

Azithromycin (AZM) is central for the treatment of chronic respiratory diseases (CRD) but has divergent off-target effects. We synthesised AZM Derivatives 1 and 2 (D1/D2) that were predicted to permit autophagy and preserve AZM’s anti-inflammatory effect. The 16HBE14o- airway epithelial cell model was exposed to AZM, D1 and D2 for 16 hr and assessed for autophagy flux via LC3B-II:p62/SQSTM1 abundance (Western blot). Necrosis was quantified via lactate dehydrogenase release. Inflammation (IL-6 secretion) was assessed in the THP-1 macrophage model exposed to 10 ng/mL lipopolysaccharide vs co-treatment with AZM and the derivatives for 18 h. AZM-derivative antibacterial activity (vs AZM) was determined via the minimum inhibition concentration (MIC) method using methicillin sensitive *Staphylococcus aureus (MSSA)*. Autophagy (LC3B-II and p62/SQSTM1 abundance) was not altered by the two derivatives and was indistinguishable from the control exposure (P> 0.05 for D1 and D2, each 10 and 50 µg/mL, vs control). D2 elicited a significant decrease in LPS-induced IL-6 secretion vs the LPS-only exposure (58.22 pg/ml, n=3, 95% ± CI [6.521 – 109.9]). Importantly, D2 caused a similar reduction in LPS-induced IL-6 secretion, as observed for AZM (-10.30 pg/mL, n=3, 95% CI [-62.00 to 41.39]). The MIC of AZM for *MSSA* growth was 0.5 µg/ml, where as D1 and D2 were 1.0 and 8.0 µg/mL, respectively (P<0.05). We show for the first time that AZM can be redesigned to mitigate its potent arrest of autophagy while preserving its anti-inflammatory activity, to counter the generation of further AZM resistant strains.

## Introduction

Chronic respiratory diseases (CRDs) put tremendous resource strain on the healthcare and economic systems. According to the Australian Institute of Health and Welfare, in 2022, approximately 2.5% of Australians suffered from chronic obstructive pulmonary disease (COPD), equating to approximately 638,000 individuals.^1^ Furthermore, within the same year, AUD$831.6 million was used to manage COPD.^1^ Globally, COPD was attributed to 3.5 million deaths (5% of deaths vs. all cause diseases) in 2021.^2^ It was estimated that COPD will incur $3.89 trillion on the global healthcare system in 2025, and by 2050, the total cost could be approximately $24.35 trillion.^3^ Diffuse panbronchiolitis, cystic fibrosis (CF), non-cystic fibrosis bronchiectasis, post-transplant bronchiolitis obliterans syndrome and chronic rhinosinusitis are amongst the other common CRDs.^4,5^

The clinical presentations of CRDs include prolonged inflammation and mucus hyperproduction, which cause obstructive airflow and bacterial colonisation.^4^ Exacerbation which is acute worsening of these conditions can occur; for example, in the case of COPD, this is indicated by ameliorated frequency of cough, dyspnoea and diminished lung function.^6^ The macrolide antibiotic azithromycin (AZM) is a frontline treatment option used for the management of these conditions - not always for its antibacterial activity, but often for patients to benefit from its rare immunomodulatory effects and its ability to accumulate intracellularly up to 100-fold compared to serum.^7,8^ In bronchiolitis obliterans, AZM is used as the first-line treatment after lung transplantation, as it is shown to improve pulmonary function by up to 18%.^7,9^ A study by Wong et al. in 2012 showed significant improvement in pulmonary function as well as a reduction in the incidence rate of exacerbations by 42% after treating patients with non-CF bronchiectasis for one year with AZM.^10^ A retrospective study in 2011 showed that administration of AZM to patients with diffuse panbronchiolitis significantly improved the symptom-burdens and sometimes enabled a curative outcome for approximately 30% of patients.^11^ Most importantly, AZM is also highly integrated in the management of COPD and CF, as it has been proven to decrease exacerbations and improve quality of life metrics.^12–15^ The British Thoracic Society has integrated AZM in its most recent guidelines for the treatment of respiratory diseases, outside its antibiotic effect to prevent exacerbations.^16^

As an antibiotic, AZM offers significant benefits compared to other antimicrobials, especially for the resolution of bacterial infections that reside in the airway. Like other macrolides, AZM exerts its antibacterial activity via binding to the polypeptide exit tunnel on the 23S rRNA near the peptidyl transferase centre (PTC) of the large ribosomal 50S subunit.^17,18^ This inhibits peptide bond formation, thereby inhibiting protein synthesis.^17,18^ The major off-target anti-inflammatory activity of AZM are characterised by the decrease of pro-inflammatory cytokines, including IL-8, IL-6, GM-CSF and TNF-α.^19,20^ AZM has also been attributed to suppression of neutrophil recruitment and degranulation.^21–23^ Further to this, AZM inhibits nuclear factor-κβ (NF- κβ), which plays an influential role in initiating and potentiating inflammatory responses.^24,25^ Moreover, AZM has been shown to polarise alveolar macrophages towards the anti-inflammatory M2 phenotype vs the pro-inflammatory M1 response, which is synonymous with CRDs.^26,27^ This effect is amplified by AZM’s capacity to enters the macrophages intracellular environment 200 times more than its levels in the extracellular matrix.^18^ Interestingly and less well described, AZM has also been shown to significantly improves epithelial barrier integrity, pointing to a reparative capacity, which is invaluable, especially for the management of CF.^28–30^

Of major concern to the development of AZM-resistance (AZM-R), current clinical practices allow AZM to be administered at a low dose that is sub-lethal to bacteria, e.g. 250 mg or 500 mg daily purely for its anti-inflammatory effect.^9,12,14,31–33^ At these doses, long-term prescription has promoted the development of clinically observable AZM-R forms of bacteria.^12,14,34–38^ In a three-year study involving low-dose administration of AZM in CF patients, after six months of treatment, all colonised *S. aureus* strains were observed to be completely resistant to macrolide antibiotics.^12^ Moreover, in the 2013 bronchiectasis intervention Study by Valery et al., who studied the effects of long-term AZM prescription vs exacerbation reduction in Indigenous children with non-CF bronchiectasis, 46% of the AZM cohort developed AZM-R.^34^ Underscoring the issues for AZM-R, antimicrobial resistance (in general) is an emerging threat to global health, and it was estimated that in 2019, 1.27 million deaths were caused by resistant species, surpassing the total mortality due to both HIV and malaria combined.^39^ AZM is currently indispensable for the effective management of CRDs. The likely emergence of AZM-R strains that have significant consequences for respiratory health will render an essential treatment mode ineffective, thereby leading to a new level of disease burden for a vast sub-set of the global population.

*In vitro* testing has also demonstrated that AZM potently inhibits autophagy macroautophagy (hereafter referred to simply as autophagy).^40–42^ Autophagy is a fundamentally important intracellular process that (amongst other functions) bridges catabolism and metabolism, by recycling redundant macromolecules and breaking down damaged organelles to release new biochemical intermediates for cellular processes.^43,44^ Critical for CRD, autophagy is also the primary intracellular bacterial clearance mechanism in mammalian cells (a process called xenophagy).^45^ As such, over-prescription of sub-antibacterial AZM can not only promote AZM-R but also arrest the very cellular process that sustains innate bacterial immunity.

AZM inhibits autophagy in a manner well known for chloroquine, called ion-trapping.^46^ AZM enters a lysosome following an ionic gradient (AZM is a unique macrolide that contains two basic amine groups) becomes protonated and overall increases the lysosomal pH (alkalinisation).^47–51^ This completely restricts lysosomal degradation by raising the pH and by preventing the activation of acid hydrolases which normally dismantles the cargos sequestered in the autophagosome.^40,46^ In pulmonary diseases, dysregulation of autophagy unto itself has been shown to contribute to the pathogenesis of several CRDs including COPD and CF.^52^ In COPD, macrophages were shown to lose the ability to carry out xenophagy, in which the fusion between the autophagosome and lysosome is disrupted.^53^ This scenario provides one explanation for the presence of persistent bacterial colonisations observed in patients.^53^ In CF, the common phenylalanine deletion mutation at position 508 of the cystic fibrosis transmembrane conductance regulator (CFTR) (CFTR^F508del^) is linked with increased production of reactive oxygen species, thus inhibiting autophagy and resulting in increased protein aggregates and potentiation of inflammation.^54^ Given downregulation of autophagy has been well documented in COPD (the most frequent and lethal CRD), this scenario for AZM where it is administered at sub-antibiotic concentrations to treat the inflammation may have unintentionally contributed to the bacteria-driven aspects of CRD and the development of AZM-R species.^55^

Given these current issues for AZM and the management of CRD, attempts have been made to modify AZM – to preserve only the immunomodulatory activity – and thereby mitigate the aforementioned consequences related to AZM-R.^30,56^ However, to date, no attempt to alter the AZM chemical structure have successfully addressed both autophagy inhibition and the retention of its anti-inflammatory effects. A derivative of AZM that is non-antibacterial but permits normal autophagy and is anti-inflammatory is in high demand and urgently needed for CRDs. This promises to address the inflammation in CRDs without further AZM-R pressure being exerted in the clinic, while permitting functional innate pathogen-removal (i.e. via xenophagy). Hence, we have synthesised derivatives 1 and 2 (D1/D2) by modifying the charged state of AZM in a manner predicted to address the above criteria. The evaluation of the AZM derivatives shows anti-inflammatory activity, modulation of antibacterial capacity and permission of autophagy activity.

## Methods

### Human Airway Epithelial Model

16HBE14o- Human Bronchial Epithelial cell line (Cat. # SCC150) was obtained from Sigma-Aldrich®. Culture media consisted of Minimum Essential Medium Eagle (Sigma-Aldrich®, M2279-500 mL), supplemented with foetal calf serum (Bovogen Biologicals Pty Ltd., 1× penicillin/streptomycin (Thermo Fisher Scientific) and 2mM L-glutamine (GlutaMAX™, Thermo Fisher Scientific). Culture conditions were 37°C in humidified atmosphere with 5% CO2. Cells were seeded at the density of 300,000 cells/mL/well, in 12-wells plates, which were subsequently left to adhere overnight prior to exposure to the respective treatment for 16 h. Treatments and their abbreviations are as follows: included no treatment (NT), AZM 10 µg/mL (AZM 10), AZM 50 µg/mL (AZM 50), Derivative 1 10 µg/mL (D1 10), Derivative 1 50 µg/mL (D1 50), Derivative 2 10 µg/mL (D2 10), Derivative 2 50 µg/mL (D2 50), Torin-1 6.1 µg/mL (Torin 6.1) (10 µM) and Chloroquine 32 µg/mL (CQ 32) (100 µM). Conditioned media samples were collected, and protein was extracted for necrosis and autophagy assessment respectively.

### Quantification of total protein

Proteins were extracted using a mixture consisting of Mammalian Protein Extraction Reagents (M-PER) (Thermo Fisher Scientific), Halt Protease and Phosphatase Inhibitor Cocktail (1:100 ratio, Thermo Fisher Scientific) and phenylmethylsulfonyl fluoride (10 µL of a 10 mM stock per millilitre of protein lysis buffer). For protein extraction and buffering, 65 µL of M-PER cell lysis buffer was added to monolayers, was scraped to collect the entire sample and then transferred to 1.5 mL Eppendorf tubes. The samples were stored in -80°C until quantified.

Total protein was quantified to enable even sample loading using the Pierce™ BCA Protein Assay Kit (Thermo Fisher Scientific). Standards were diluted from 2 mg/mL to 0.0625 mg/mL with 4 replicates of 10 µL per well for each concentration. Experimental samples were similarly prepared in replicates of three (10 µL each well). Colourimetry detection was performed by mixing Reagent A and B in 1:50 ratio, introduced to each well 200 µL, mixed with samples or standards, incubated in 37°C for 25 min and read at 562 nm. Sample protein concentration was determined for each sample using the standard curve method to enable 10 µg of protein were determined.

### Western blot

Electrophoresis was performed at 100V for approximately 2 h, using 15-wells NuPAGE™ 4-12% Bis-Tris Gel (Thermo Fisher Scientific) in Bolt™ MOPS SDS running buffer (Novex® by Life Technologies). Each well of the gel was loaded with 15 µL of a mixture of MiliQ water, 10 µg of sample, 3.75 µL of Loading Buffer (Thermo Fisher Scientific) and 1.5 µL Reducing Agent (Thermo Fisher Scientific). Proteins were then transferred to Nitrocellulose Mini Format membrane (Bio-Rad) at 1.3 A, 2.5 V for 7 min. The membranes were sectioned into three distinct sections corresponding to the protein’s molecular weights consistent with ß-actin, LC3B-I/II and p62/SQSTM1/Sequestosome 1 (for autophagy assessment).

Skim milk 5% and Bovine Serum Albumin 5% (Sigma-Aldrich) were used for blocking of ß-actin, and LC3B-I/II and p62/SQSTM1 blots respectively for 1 h. Mouse anti-ß-actin (1:10,000, Cell Signalling Technology, A1978), rabbit anti-p62/SQSTM1 (1:3000, Cell Signalling Technology, 5114S) and rabbit anti-LC3B-I/II (1:3000, Cell Signalling Technology, 4108S) were used for primary incubation. Primary antibody incubation was conducted in 4°C condition, overnight with 40 rpm shaking. The membranes were washed with TBS-T four 15 min intervals. Anti-mouse (1:10,0000, RD System®) and anti-rabbit (1:4000, secondary HRP conjugated IgG) were suspended in 5% skim milk in TBS-T for the secondary antibody incubation. Secondary antibody incubation lasted for one hour followed by the four TBS-T washes.

Blots were imaged by applying Novex™ ECL Chemiluminescent Substrate Reagent (Invitrogen™) in 1:1 ratio of Reagent A and Reagent B 5 min before being imaged; for ß-actin blots, no incubation period was conducted. Images were acquired using the Fujifilm® LAS-4000 platform, and quantification was performed using MultiGauge V3.0 software (Fujifilm). Band intensities of all proteins were quantified (and background signal subtracted) and normalised the β-actin loading control to determine a final signal magnitude. Data was recorded and analysed in Excel and GraphPad Prism 10.6.1.

### Necrosis Assessment Assay

To assess any potential necrotic influence of the exposures, lactate dehydrogenase (LDH) released assay was performed in conditioned media entered into the CyQUANT™ LDH Cytotoxicity Assay (Cat. # C20300 and C20301).

The reaction mixture comprised of substrate stock solution and assay buffer (1:200 ration mix). Firstly, the substrate stock solution was made by reconstituting the Substrate Mix in 11.4 mL of distilled water (Milli-Q). Then, 600 µL of assay buffer was added to complete the reaction mixture. In each well of a 96-well plate, 50 µL of Reaction Mixture was added to 50 µL of treatment media. The plate was left for incubation for 30 min in a dark environment. Lastly, 50 µL of Stop Solution was added and a readout was performed at 490 nm and 680 nm. Analysis of data was carried out in GraphPad Prism, by subtracting the reading at 680 nm from 490 nm. All readings were normalised to the positive control (maximum LDH released) prepared by exposing cells to 2% Triton-X.

### Minimum Inhibition Concentration Assessment of Antibiotic Activity

Minimum inhibition concentration (MIC) assays were performed to determine the antibiotic activity of the AZM derivatives vs AZM. The samples were tested against methicillin-sensitive *Staphylococcus aureus (MSSA)* ATCC 25923, following the European Committee on Antimicrobial Susceptibility Testing (EUCAST) in ISO 20776-1:2019.^57^ Data were plotted in Excel and GraphPad Prism 10.6.1. Two-way ANOVA was used to compare the significance between the derivatives and the parent AZM. The agents tested included azithromycin (AZM), Derivative 1 (D1), Derivative 2 (D2) and azithromycin oxime (AZM-OX).

The standard medium used was cation-adjusted Mueller-Hinton broth (caMHB). The tested concentration range was 64 to 0.125 µg/mL, achieved by two-fold serial dilution. Preparation of inoculum comprised of suspending a fresh MSSA colony in sterile saline (0.9% w/v). A densitometer was used to adjust the turbidity to 0.5 McFarland standard.

96-Well polystyrene microplates from Corning® with flat bottom were prepared with 100 µL of caMHB broth in all wells, except for wells of column 1, which held 200 µL each well. Each compound was pipetted into two adjacent wells within a column, starting from column 1 as technical repeats, and at twice the intended concentration. From column 1, two-fold serial dilutions were performed using 100 µL of the antibiotic-broth mixture up to column 10, at which 100 µL was discarded from each well. Positive and negative (sterile) controls were designated for columns 11 and 12, respectively. All wells of columns 1 to 11 were then added with 100 µL of the inoculum, equivalent to a final density of 5 × x105 CFU/mL, to achieve the intended testing concentrations.

Optical density reading at 595 nm (OD595) on day 0 was taken with Thermo Scientific Multiskan FC plate reader to generate baseline reading. The plates were then incubated for 16 h h at 37°C on an orbital shaker set at 200 rpm and subsequently read at 595 nm. MIC was defined as the lowest concentration of the compound in which the change in OD595 was less than 0.1.

### Assay and Model of Pro-inflammatory IL-6 Secretion

The THP-1 macrophage model was used to examine a system that produces a strong inflammatory response (here IL-6 secretion) due to stimulation with lipopolysaccharide (LPS). The Human Acute Monocytic Leukaemia Cell Line (THP-1; purchased from the American Type Culture Collection, ATCC-TIB-202™) were propagated in RPMI 1640 culture media (Thermo Fisher Scientific) was supplemented with foetal bovine serum, penicillin/streptomycin and L-glutamine according to previously stated methods, with the addition of 2-mercaptoethanol (Sigma-Aldrich) to 50 µM. For differentiation, the cells were seeded at 300,000 cells/mL/well and treated with phorbol 12-myristate 13-acetate (PMA) at 50 ng/mL. The cells were incubated in PMA media for 24 h, and then the media was changed for standard RPMI to allow a 48-h rest period. Differentiation was qualified by THP-1 cells (normally non-adherent) attachment to the culture well growth surface. Exposure to respective AZM/AZM-derivative (and controls) conditions were for 18 h. The media was collected for analysis using Human IL-6 ELISA kit (88-7066, by Thermo Fisher Scientific). Outcomes were for n = 3 repeats cultures.

### IL-6 ELISA Protocol

Coating antibody was diluted 1:250 in 1× DPBS and incubated overnight (100 µL per well) and then washed with PBS-T (0.05% Tween-20) for three times. 1× ELISA diluent was prepared by diluting ELISA Diluent concentrate in 1:5 ratio with MilliQ water. 200 µL was pipetted into each well and the plate was incubated for 1 h at room temperature. The plates were then washed twice with PBS-T. An 8-point standard curve was prepared by performing a two-fold serial dilution of IL-6 standard reconstituted with MilliQ water (n=2 technical replicates per concentration) starting at 200 pg/mL. Samples were added for n=3 technical replicates per sample) with 100 µL per well. The plate was incubated overnight in 4°C then washed four times with PBS-T. Followed was the addition of 100 µL 1× detection antibody (diluted in 1x ELISA diluent). The plate was sealed and incubated for 1 h at room temperature and unbound substrate removed via four times of PBS-T washes. Horseradish

Peroxidase (HRP) was diluted 1:100 in 1× ELISA diluent and 100 µL was pipetted into each well. The plate was further incubated for one hour, followed by six PBS-T washes. 100 µL of 1× TMB was added to each well and incubated for 15 min at room temperature. 1 N H2SO4 was added at a volume of 100 µL to each well (stop solution) and then the plate was read at 450 nm.

### Statistical analysis

Data were recorded in Microsoft Excel and later transferred to GraphPad Prism 10.6.1 for analysis. One-way ANOVA was conducted for autophagy (Western blot), necrosis (LDH) and anti-inflammatory (ELISA) assessment. Antibacterial (MIC) analysis was performed on two-way ANOVA basis.

## Results

### Human Airway Epithelial Cells Exhibit Normal Viability When Exposed to AZM and the Derivatives

First, we tested whether the derivatives induce cytotoxicity in airway epithelial cells. Upon microscopic observation, there was no morphological difference between the untreated cells and the derivatives-exposed counterparts at 50 µg/mL (**Fig 1A-C**). Cells undergoing necrosis will release lactate dehydrogenase (LDH), a cytoplasmic enzyme that converts NADH to NAD^+^ and pyruvate to L-lactate.^58^ As LDH is also stable in media, it serves as a marker of compromised cells when extracellular and indicates necrosis. Similarly, LDH assessment showed no significant difference in cell necrosis between treatments (p>0.05, **Fig 1D**), indicating that the derivatives were safe as per AZM. Of note on cell toxicity, BCA protein quantification showed similar protein levels across all samples, indicating normal cell functions (data not shown). In reference to the Western blots, similar β-actin bands were observed across all exposures, demonstrating viable cells.

**Figure 1:**
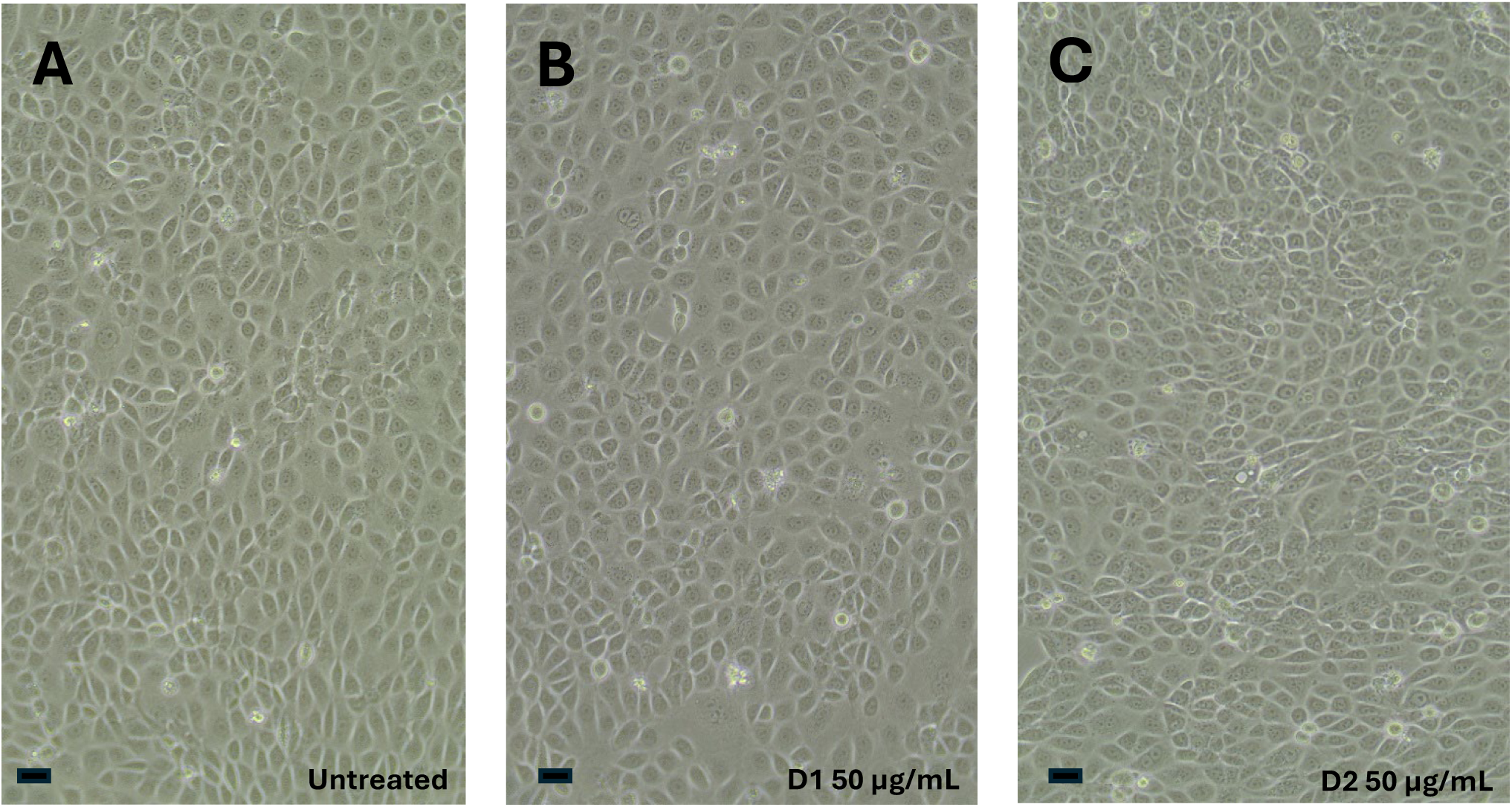
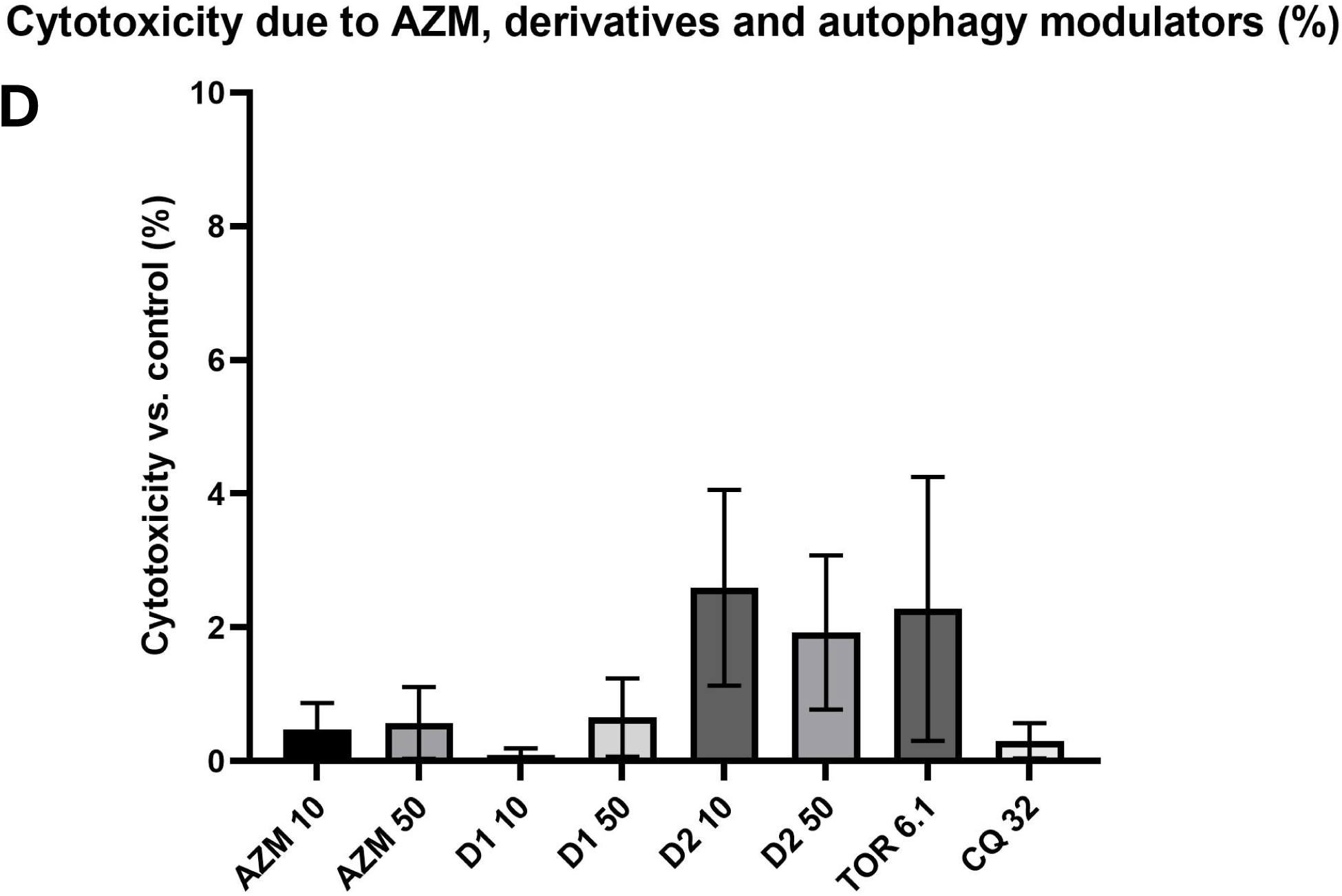
The AZM derivatives exhibit a safe toxicity profile when exposed to human airway cells. 16HBE14o- cells were exposed to AZM, the derivatives and autophagy modulators for 16 h. **A, B and C.** Images of 16HBE14o- cells untreated, exposed to D1 and D2 at 50 µg/mL (respectively), show similar morphology across conditions (scale bar is 50 µm). **D.** Similarly there was no evidence that the derivatives elicited LDH release (due to loss of plasma membrane integrity) as a readout of toxicity vs the parent molecule. *The data represented n = 3 independent experimental replicates. AZM 10 = AZM 10 µg/mL; AZM 50 = AZM 50 µg/mL; D1 10 = Derivative 1 10 µg/mL; D1 50 = Derivative 50 µg/mL; D2 10 = Derivative 2 10 µg/mL; D2 50 = Derivative 2 50 µg/mL; CAL = Calibrator; Torin 6.1 = Torin-1 6.1 µg/mL; CQ 32 = Chloroquine 32 µg/mL. Statistical analysis was a one-way ANOVA*.

### The Derivatives Permit Normal Autophagy

We next assessed the derivative’s influence on autophagy. D1 and D2 both alleviated the inhibition, as shown by the LC3B-II magnitude for these two agents at 50 µg/mL, that were appreciably lower vs AZM at the same concentration (6-fold and 12-fold respectively; *x_d_* = -2.95, 95% CI [-4.25 to - 1.64]; and *x_d_*= -3.19, 95% CI [-4.49 to -1.88], respectively, p<0.0001) (**Fig 2A**). This is underscored by no significant difference between D1/2 vs the ‘no treatment’ control (*x_d_* =+0.34, 95% CI [-0.96 to 1.65] and *x_d_* = +0.027, 95% CI [-1.2 to 1.41], respectively, p>0.05) (**Fig 2A**). Furthermore, the quantified LC3B-II signals of AZM 50 µg/mL were 2-fold higher than that of CQ at 32 µg/mL (positive control) (*x_d_*= +2.06, 95% CI [0.76 to 3.36], p=0.0008), is evidence that AZM blocks autophagy to a greater extent than chloroquine (**Fig 2A**). There was a non-significant trend of increasing accumulation of p62/SQSTM1/SQSTM-1 with elevating AZM concentration (**Fig 2B**).

**Figure 2:**
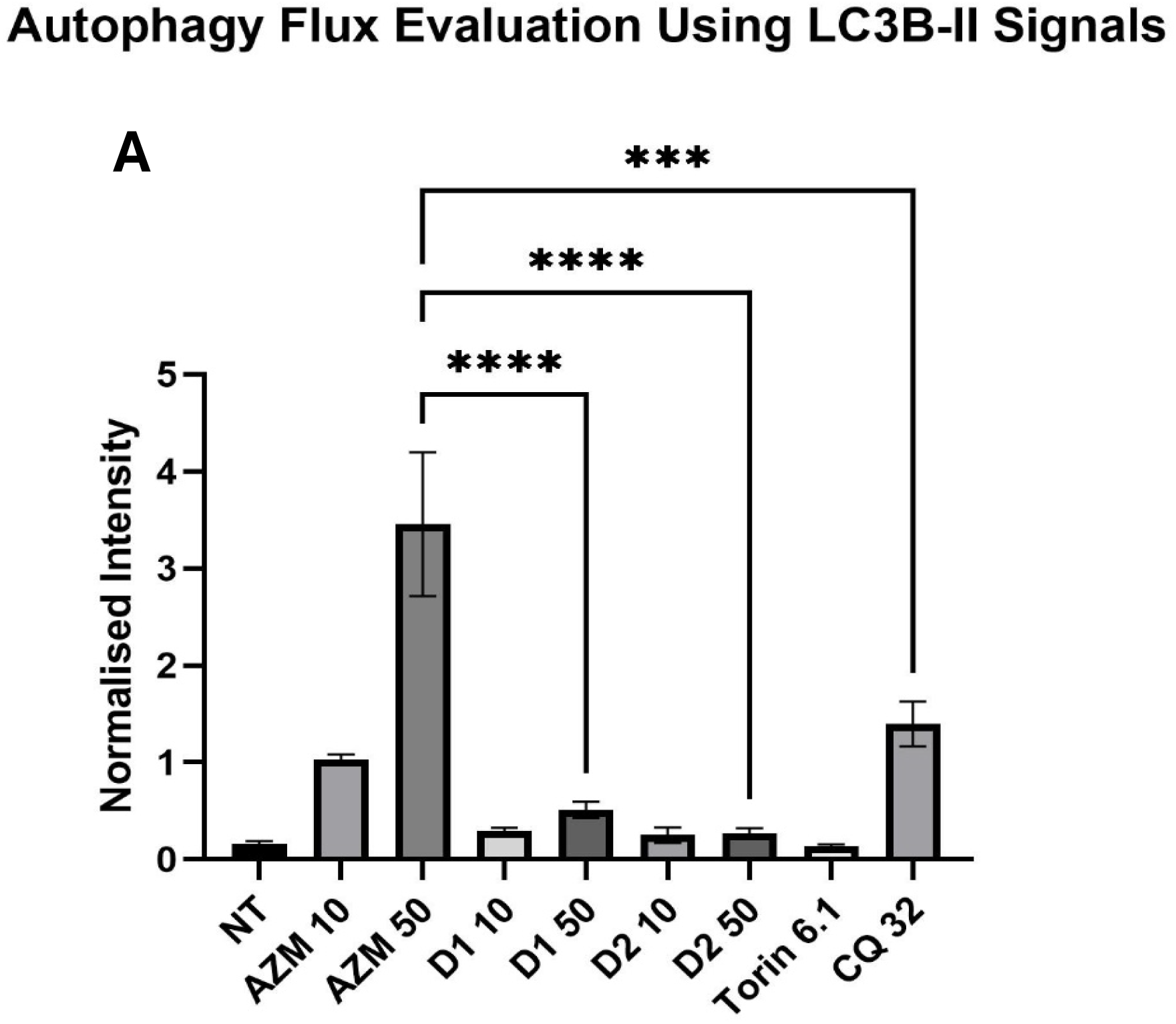
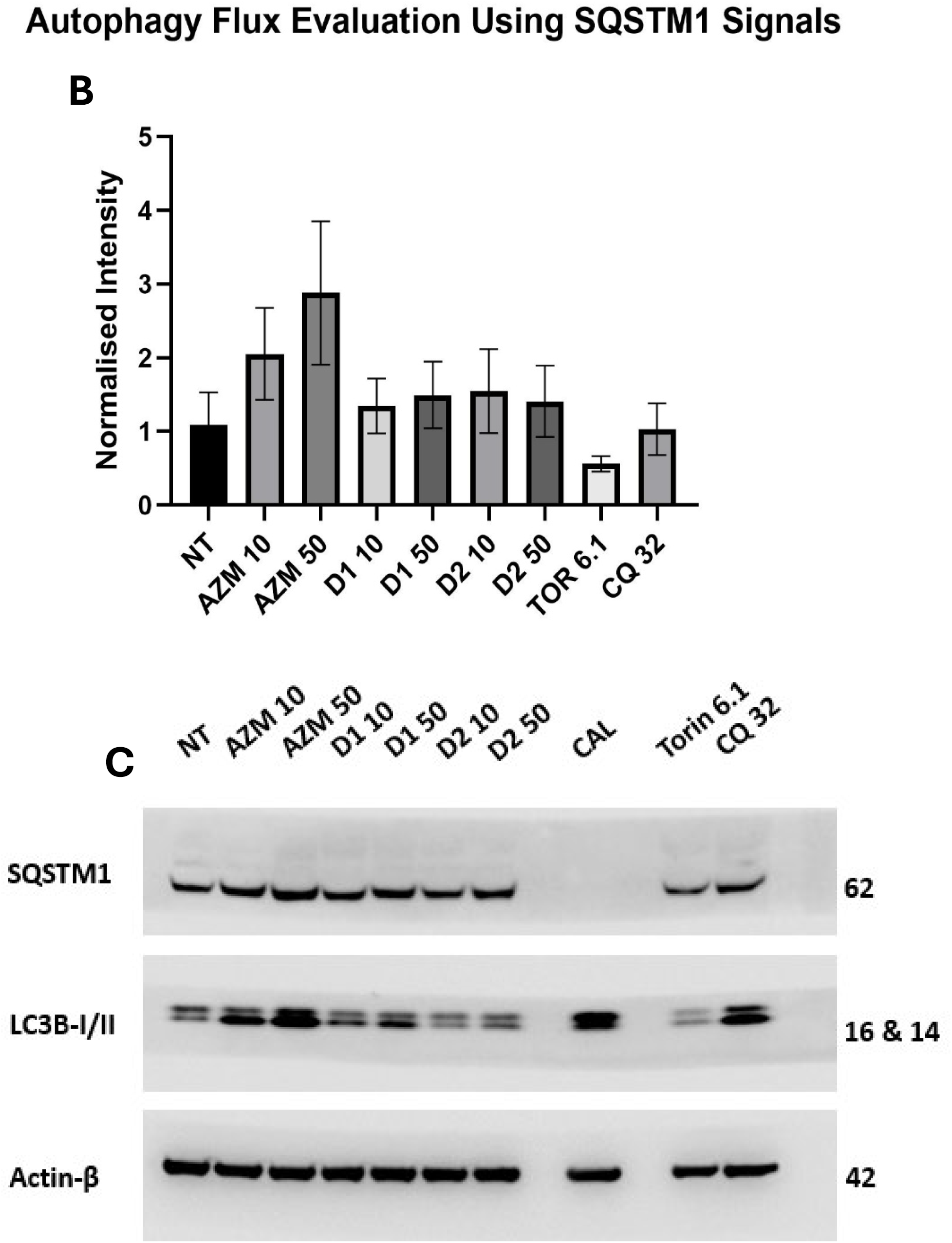
The AZM derivatives mitigate the effects of the parent molecule on autophagy. Assessment of autophagy of 16HBE14o- cells exposed to AZM, D1 and D2 and autophagy modulators torin-1 (promoter) and chloroquine (CQ; inhibitor) as controls for 16 h via Western blot. **A.** LC3B-II is elevated with AZM vs D1/D2. **B.** Quantification of P62/SQSTM1/SQSTM1 abundance shows a trend for its accumulation with AZM, while D1/D2 were similar to the NT exposure. **C.** A representative blot confirming the modulation of autophagy in 16HBE14o- cells for LC3B-II and P62/SQSTM1/SQSTM1 (normaliser was ß-actin). *NT = no treatment; AZM 10 = AZM 10 µg/mL; AZM 50 = AZM 50 µg/mL; D1 10 = Derivative 1 10 µg/mL; D1 50 = Derivative 50 µg/mL; D2 10 = Derivative 2 10 µg/mL; D2 50 = Derivative 2 50 µg/mL; CAL = Calibrator; Torin 6.1 = Torin-1 6.1 µg/mL; CQ 32 = Chloroquine 32 µg/mL. Quantified band intensity for each treatment of each protein was normalised against the corresponding ß-actin reading. Outcomes are representative of n=3 independent measure. One-way ANOVA statistical analysis was used, and data are presented as mean±SEM.; P < 0.001 ***, P < 0.0001 ****. Statistical analysis was one-way ANOVA*.

While p62/SQSTM1 signals across treatments remained statistically stable (p>0.05), this concomitant with elevated LC3B-II qualifies the findings for autophagy. Regardless, the modification of AZM in its charged state has been shown to mitigate the block on autophagy. It is noteworthy that the outcomes for the derivatives (i.e. enabling normal autophagy) were also recapitulated in the THP-1 model used to assess immunomodulation (data not shown).

### Derivative 1 Retained the Antimicrobial Effects Observed in AZM

After we had assessed autophagy, we wanted to determine if the modifications also affected their antimicrobial activity. MSSA growth profiles showed that D1 was similarly mitostatic to bacterial growth as the parent molecule (zero detectable growth at 1.0 µg/mL). In contrast D2 exhibited (approximately 8-fold) diminished antimicrobial activity. AZM oxime (AZM-OX) was included as an example of our previously synthesised AZM derivative that also enabled autophagy, but that removed all antimicrobial activity (**Fig 3**).^55^ Hence, this presents interesting structure-activity relations information as this relates to these modifications.

**Figure 3:**
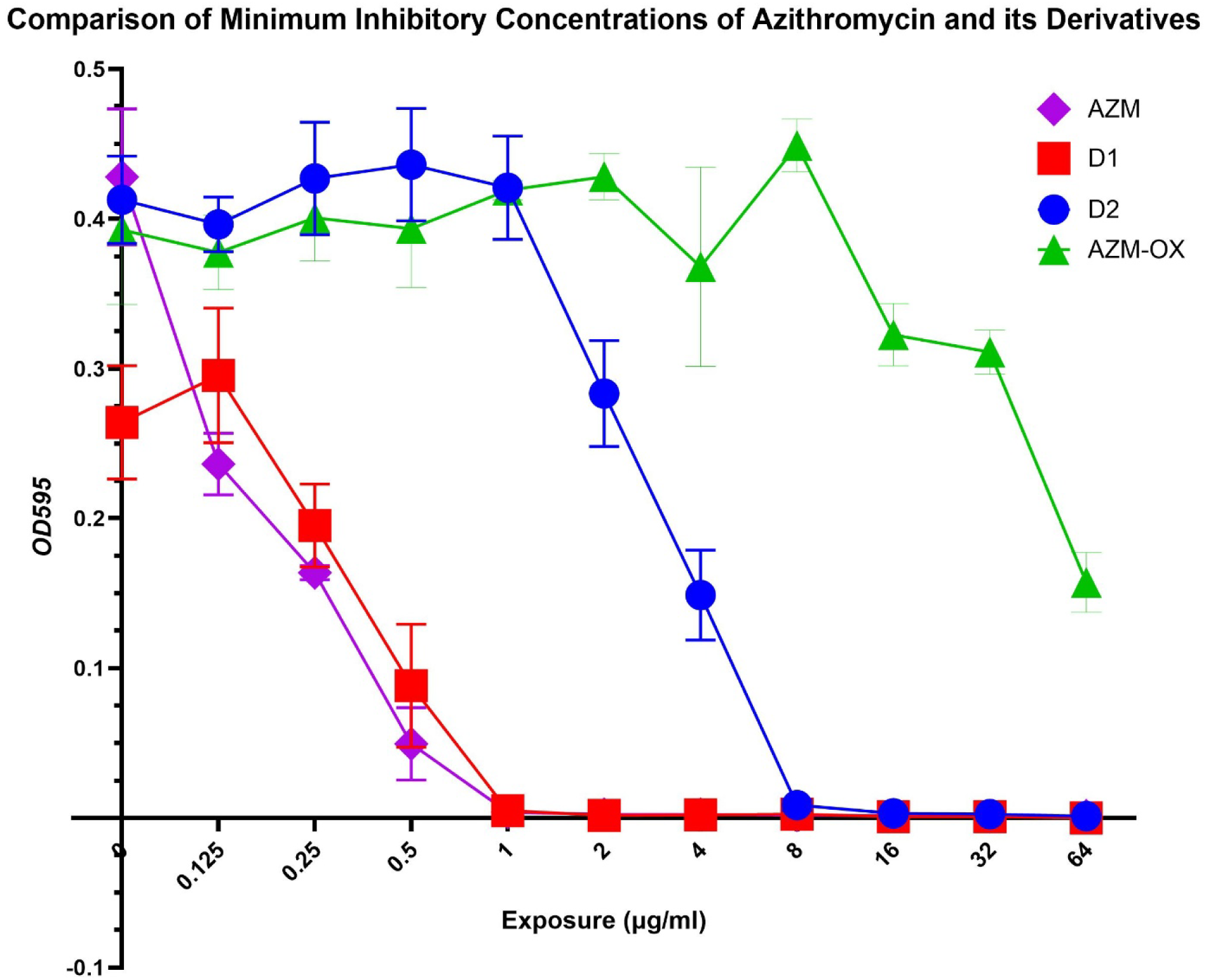
Alterations to the derivatives produced differential antimicrobial outcomes. MIC was used to evaluate the antibiotic potential of AZM, D1, D2 and AZM-oxime (AZM-OX) in a methicillin susceptible MSSA model. *Data is representative n=4 independent experiments. Data are mean±SEM. Statistical analysis was two-way ANOVA*.

### Derivative 2 is Anti-inflammatory

We next assessed the immunomodulatory capacity of the derivatives. Importantly, D1, D2 and AZM were not pro-inflammatory (*x_d_* = -3.04, 95% CI [-54.73 to 48.66]; *x_d_* = -3.51, 95% CI [-55.21 to 48.18]; and *x_d_* = -2.39, 95% CI [-54.09 to 49.30], respectively, p>0.05 (**Fig 4**). Administration of AZM to LPS-stimulated THP-1 macrophages decreased the secretion of IL6 significantly (*x_d_* = - 68.52, 95% CI [-120.2 to -16.83], p = 0.0018). Immunomodulatory capability was not observed in D1 (*x_d_* = -4.6, 95% CI [-56.29 to 47.10], p>0.05. On the other hand, D2 retained the anti-inflammatory activity as observed in AZM (*x_d_* = -58.22, 95% CI [-109.9 to -6.52], p = 0.0153), at a similar magnitude as the parent molecule (*x_d_* = +10.30, 95% CI [-41.39 to 62.00], p>0.05).

**Figure 4:**
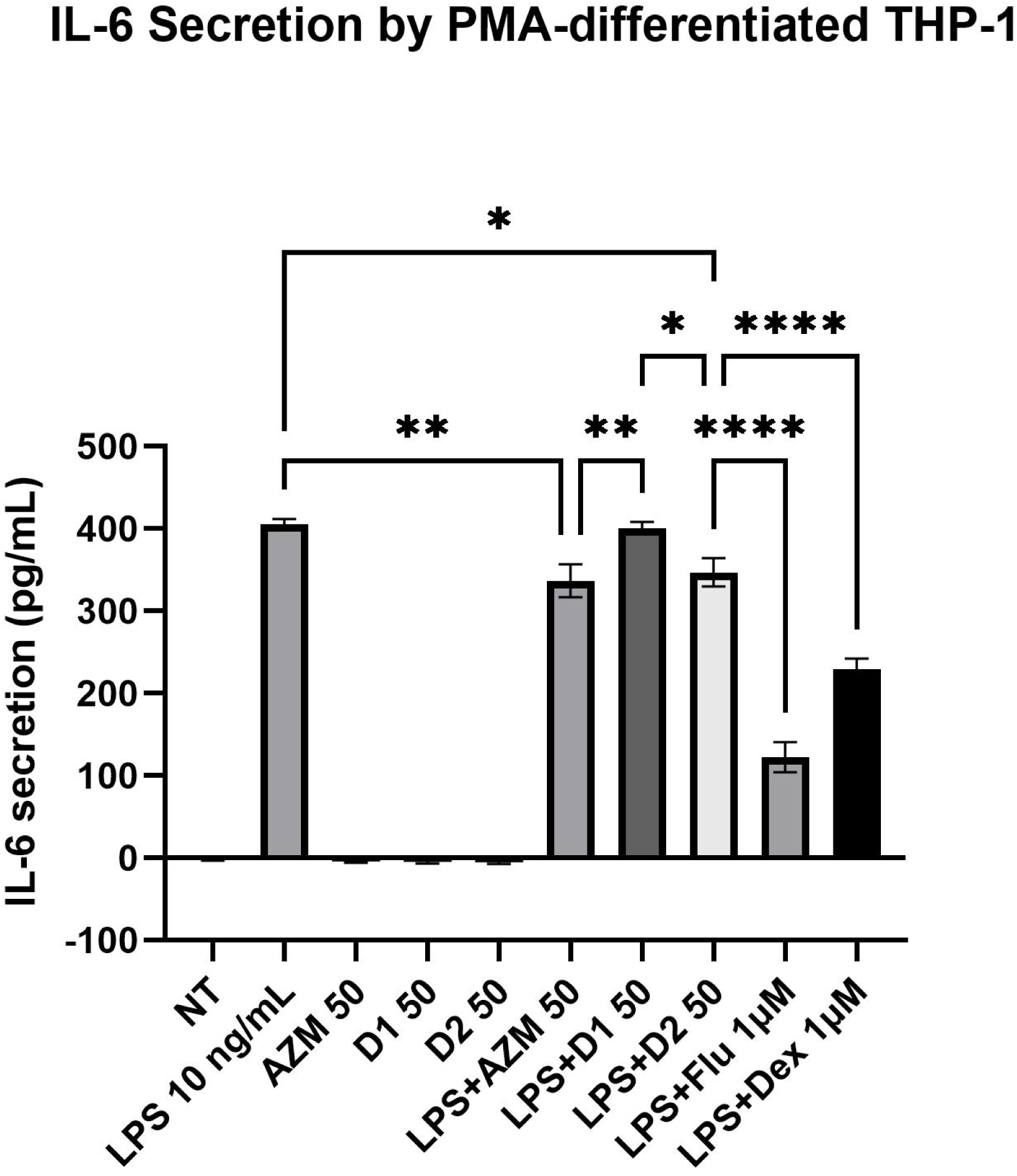
Derivative 2 exhibits immunomodulatory activity consistent with AZM. IL-6 secretion in the PMA-differentiated THP-1 macrophage model was determined using ELISA. THP-1 cells were differentiated in 50 ng/mL PMA for 24 h. The macrophages were then exposed to LPS 10 ng/mL and co-treated with AZM (AZM), Derivative 1 (D1), Derivative 2 (D2), fluticasone (Flu) and dexamethasone (Dex) for 18 h. *Dexamethasone/fluticasone were used as clinically relevant positive control for LPS-induced IL-6 suppression. Unless stated, concentrations are µg/mL. Data is representative of n=3 independent experiments. Data presented as mean±SEM. All data was analysed with one-way ANOVA. * P < 0.05, ** P < 0.01, **** P < 0.0001*.

## Discussion

The antimicrobial and anti-inflammatory properties of AZM distinguishes it for CRD. Although it offers effective reduction of inflammation-related exacerbations, a major disadvantage is inhibiting autophagy in host cells. The deceleration in antibiotic innovation in the last 30 years, coupled to the increase in resistant species, means safeguarding AZM is imperative.^59^ Herein we described AZM derivatives that permitted normal autophagy, exhibited anti-inflammatory activity and retain antibacterial effect.

Autophagy is the most effective intracellular bacterial clearance mechanism. D1 and D2 enable autophagy at levels consistent with normal cell function. This was evidenced by the release of accumulated autophagy proteins (LC3B-II and p62/SQSTM1) vs AZM.^40,41,55,60^ Interestingly, AZM was a more influential inhibitor of autophagy than CQ, a commonly employed negative control.

Indeed, this occurred at a lower concentration of AZM vs CQ (66.76 µM vs 100 µM, respectively; **Fig 2A**). This is supported by in Takano et al. 2023 who applied CQ and AZM (both 50 µM) and assessed autophagy using a viral reported system.^61^ Both D1 and D2 showed significant decreases in autophagy inhibition at both 10 and 50 µg/mL compared to AZM (6-fold and 12-fold, respectively). While this was effectively exhibited for LC3B-II abundance, p62/SQSTM1 accumulation was noted only as a trend (vs statistically significant). This is likely due to already high baseline levels of p62/SQSTM1 for the NT conditions. This outcome for LC3B-II and p62/SQSTM1 as markers for autophagy was also noted previously for the 16HBE14o- model.^55^ Nevertheless, and taken together, these outcomes show that the modifications applied to D1/D2 generated AZM species that permit host cell bacterial clearance.

The most sought after outcomes when generating AZM derivatives are: 1. the clearance of AZM-R bacteria, or 2. syntheses that lack antibiotic activity – i.e. which mitigate AZM-R but impart the anti-inflammatory effects. Regarding enhanced antimicrobial effect on AZM-R species, Zhang and colleagues (2010) synthesised 3-*O*-carbonyl derivatives of 11,12-cyclic carbonate AZM by introducing extended 3-*O*-carbamoyl side chains.^62^ While select derivatives showed significant activity against erythromycin-resistant *S. pneumoniae* with *erm* and *mef* genes (AB11),^62^ none of the 11,12-cyclic carbonates derivative exhibited effects against *S. aureus* ATCC 25923.^62^ The 4’’-*O*-(trans-ß-arylacrylamido)carbamoyl analogues of AZM were synthesised by incorporating an arylacrylamido group into the 4’’-position.^63^ While some analogues showed MIC values of 0.5 µg/mL against *S. aureus* ATCC 25923, others were more effective at inhibiting *S. pneumoniae* AB11 at 1 µg/mL, 256-fold more than AZM.^63^ Our D1 also exhibited antibacterial effect comparable to the original AZM. However, the notable advantage that D1 has over AZM and other derivatives is the permission of normal autophagy. Hence, our AZM derivatives show great promise to resolve inflammation for individuals suffering from CRD *and* permit both host-cell bacterial immunity (i.e. autophagy) and the antibiotic effect elicited by this next-generation AZM. D1 also provides an opportunity to investigate structure–activity relationships underlying AZM-associated autophagy inhibition, as it retains antibacterial activity comparable to AZM while substantially reducing its effects on autophagy-associated markers. Albeit having a diminished antibacterial effect compared to AZM and D1, D2 offers tremendous benefits to counter AZM-R. By exerting less selective pressure, bacteria are less likely to develop tolerance to D2. A recent study in 2021 identified that AZM was only beneficial in reducing COPD-exacerbations after a year of administration; an outcome that was attributed to the emergence of resistant species.^64^ Moreover, given the well-established link between autophagic dysregulation and CRD, further arrest of autophagy by AZM would amplify this phenomenon. It follows that, given AZM inhibits autophagy, attenuates inflammation and promotes AZM-R, these drive bacterial colonisation, thereby increasing exacerbations.^6,64^ Hence, AZM derivatives that are anti-inflammatory but lack antibacterial activity has extreme clinical value, which is underscored by AZM being primarily prescribed for its immunomodulatory effect in CRDs. While D1 completely lost its ability to attenuate inflammation, D2 significantly attenuate the inflammatory response to the same extent as AZM. With a reduced antibiotic effect and preserved immunomodulatory activity, D2 can be potentially administered at the same dose as AZM, with significantly less selection pressure against AZM. Compared to D2, AZM-OX, (the only other known AZM derivative that mitigates autophagy inhibition) did not exhibit anti-inflammatory activity, offering tremendous benefit.^55^ Another AZM derivative known as 2′-Dehydroxy-5″-Epi-Azithromycin, showed no antibacterial properties, while retaining the full immunomodulatory capabilities.^65^ The analogue CSY0073 synthesised by Synovo, also showed no antibacterial activity against common airway-colonised bacteria including *S. aureus, P. aeruginosa* and *H. influenzae.*^66^ However, it retained the anti-inflammatory capability tested in mice macrophages isolated from bronchoalveolar lavage fluids (BAL) exposed to 1 µg/mL LPS.^66^ CSY0073 decreased significantly the amount of secreted IL-6 at all concentrations tested (10, 20 and 50 µM.^66^) In comparison to previous analogues, both D1 and D2 offer compelling advantages, in which bacterial clearance capability can be enhanced, and effective management of inflammation with minimal risk of antibiotic resistance development.

Out of the two derivatives, D2 with diminished antibacterial activity and retaining the full anti-inflammatory effects seen in the native AZM molecule has high value. Although D2 is not completely non-antibiotic, it is the only derivative to date that offers these attributes while permitting normal autophagy flux. The other autophagy-permissive derivative (AZM-OX) does not preserve either of these properties.^55^ This highlights D2 as a standout candidate as a next generation AZM analogue - it allows host-cell immunity to remain functional, decreases the overall selective pressure for AZM resistance and retains clinical value as an anti-inflammatory. To ensure that D2 is eligible to become a new drug candidate, further assessments must be undertaken, including performing MIC on other bacteria species, using other cell models that resemble CRD pathophysiology e.g. air-liquid interface of primary airway models from control/COPD and peripheral blood mononuclear cell (PBMC) models. THP-1 (used here) vs PBMCs have distinct cytokine secretion profiles when exposing to pyrogens (i.e. PBMC-macrophages secrete higher levels of e.g. IL-6, IL-8 and TNF-α^67–69^).

While these findings support the potential of D1 and D2 as improved AZM analogues, they are early-stage characterisations. Progressing therapeutic development would require a comprehensive toxicity and safety evaluation consistent with pre-clinical drug pipelines.^70,71^ This would include expanded cytotoxicity screening across multiple primary cell models, assessment of apoptosis and stress pathways (e.g., caspase activation and mitochondrial depolarisation), and longer-term viability assessments, as reviewed in Tang and More, 2023.^72^ Standard profiling such as dose/response curves, maximum tolerated concentrations, and off-target activity panels would also be required. In addition, preliminary stability studies, along with antibacterial spectrum testing beyond *MSSA*, are needed for candidate development. These steps would provide the depth of safety and functional data expected by industry groups and IP evaluators.

## Conclusion

CRDs cause an immense burden to patients, and the inevitable emergence of antibiotic-resistant species, pose substantial challenges to the healthcare system. AZM, a macrolide antibiotic often prescribed off label for anti-inflammatory purposes (at sub-inhibitory doses), is of no exception. AZM is also found to inhibit autophagy, further potentiating bacterial colonisation of the airway. We developed, for the first time, a derivative of AZM that retains immunomodulatory activity while exhibiting diminished antimicrobial activity (Derivative 2). Importantly, the derivative permits normal autophagy in cells, allowing the natural intracellular bacterial clearance machinery to operate. These attributes put the derivative in a leveraged position relative to other (described) analogues.

More important, this study promises new options for the clinic and patients suffering from long-term disease and can safeguard the longevity of the parent molecule.

## Declarations

The authors have no conflicts of interest to declare.

## Acknowledgements

Samyuktha Kothandaraman for assistance with ELISA experiments.

## Reference

1. Health AIo & Welfare, Chronic obstructive pulmonary disease. 2024, AIHW: Canberra.

2. World Health Organization (2024). Chronic obstructive pulmonary disease (COPD). https://www.who.int/news-room/fact-sheets/detail/chronic-obstructive-pulmonary-disease-(copd).

3. Boers E, Allen A, Barrett M, Benjafield AV, Rice MB, Wedzicha JA, Kaye L, Zar HJ, Sinha S, Ozoh O, Crotty Alexander LE & Malhotra A (2025). Forecasting the Global Economic and Health Burden of COPD From 2025 Through 2050. CHEST 168, 880–889.

4. Spagnolo P, Fabbri LM & Bush A (2012). Long-term macrolide treatment for chronic respiratory disease. European Respiratory Journal 42, 239–251.

5. Crosbie PAJ & Woodhead MA (2008). Long-term macrolide therapy in chronic inflammatory airway diseases. European Respiratory Journal 33, 171–181.

6. Wedzicha JA & Seemungal TA (2007). COPD exacerbations: defining their cause and prevention. Lancet 370, 786–96.

7. Welte T (2019). Azithromycin: The Holy Grail to Prevent Exacerbations in Chronic Respiratory Disease? Am J Respir Crit Care Med 200, 269–270.

8. McMullan BJ & Mostaghim M (2015). Prescribing azithromycin. Aust Prescr 38, 87–9.

9. Gottlieb J, Szangolies J, Koehnlein T, Golpon H, Simon A & Welte T (2008). Long-Term Azithromycin for Bronchiolitis Obliterans Syndrome After Lung Transplantation. Transplantation 85, 36–41.

10. Wong C, Jayaram L, Karalus N, Eaton T, Tong C, Hockey H, Milne D, Fergusson W, Tuffery C, Sexton P, Storey L & Ashton T (2012). Azithromycin for prevention of exacerbations in non-cystic fibrosis bronchiectasis (EMBRACE): a randomised, double-blind, placebo-controlled trial. The Lancet 380, 660–667.

11. Li H, Zhou Y, Fan F, Zhang Y, Li X, Yu H, Zhao L, Yi X, He G, Fujita J & Jiang D (2011). Effect of Azithromycin on Patients with Diffuse Panbronchiolitis: Retrospective Study of 51 Cases. Internal Medicine 50, 1663–1669.

12. Samson C, Tamalet A, Thien HV, Taytard J, Perisson C, Nathan N, Clement A, Boelle P-Y & Corvol H (2016). Long-term effects of azithromycin in patients with cystic fibrosis. Respiratory Medicine 117, 1–6.

13. Clement A, Tamalet A, Leroux E, Ravilly S, Fauroux B & Jais JP (2006). Long term effects of azithromycin in patients with cystic fibrosis: A double blind, placebo controlled trial. Thorax 61, 895–902.

14. Albert RK, Connett J, Bailey WC, Casaburi R, Cooper JAD, Criner GJ, Curtis JL, Dransfield MT, Han MK, Lazarus SC, Make B, Marchetti N, Martinez FJ, Madinger NE, McEvoy C, Niewoehner DE, Porsasz J, Price CS, Reilly J, Scanlon PD, Sciurba FC, Scharf SM, Washko GR, Woodruff PG & Anthonisen NR (2011). Azithromycin for Prevention of Exacerbations of COPD. New England Journal of Medicine 365, 689–698.

15. Uzun S, Djamin RS, Kluytmans JAJW, Mulder PGH, van’t Veer NE, Ermens AAM, Pelle AJ, Hoogsteden HC, Aerts JGJV & van der Eerden MM (2014). Azithromycin maintenance treatment in patients with frequent exacerbations of chronic obstructive pulmonary disease (COLUMBUS): a randomised, double-blind, placebo-controlled trial. The Lancet Respiratory Medicine 2, 361–368.

16. Smith D, Du Rand I, Addy CL, Collyns T, Hart SP, Mitchelmore PJ, Rahman NM & Saggu R (2020). British Thoracic Society guideline for the use of long-term macrolides in adults with respiratory disease. Thorax 75, 370–404.

17. Schlünzen F, Harms JM, Franceschi F, Hansen HAS, Bartels H, Zarivach R & Yonath A (2003). Structural Basis for the Antibiotic Activity of Ketolides and Azalides. Structure 11, 329–338.

18. Parnham MJ, Haber VE, Giamarellos-Bourboulis EJ, Perletti G, Verleden GM & Vos R (2014). Azithromycin: Mechanisms of action and their relevance for clinical applications. Pharmacology & Therapeutics 143, 225–245.

19. Cramer CL, Patterson A, Alchakaki A & Soubani AO (2017). Immunomodulatory indications of azithromycin in respiratory disease: a concise review for the clinician. Postgraduate Medicine 129, 493–499.

20. Yang J (2020). Mechanism of azithromycin in airway diseases. Journal of International Medical Research 48, 0300060520932104.

21. Tsai WC, Rodriguez ML, Young KS, Deng JC, Thannickal VJ, Tateda K, Hershenson MB & Standiford TJ (2004). Azithromycin Blocks Neutrophil Recruitment in Pseudomonas Endobronchial Infection. American Journal of Respiratory and Critical Care Medicine 170, 1331–1339.

22. Èulić O, Eraković V, Čepelak I, Barišić K, Brajša K, Ferenčić Ž, Galović R, Glojnarić I, Manojlović Z, Munić V, Novak-Mirčetić R, Pavičić-Beljak V, Sučić M, Veljača M, Žanić-Grubišić T & Parnham MJ (2002). Azithromycin modulates neutrophil function and circulating inflammatory mediators in healthy human subjects. European Journal of Pharmacology 450, 277–289.

23. Verleden GM, Vanaudenaerde BM, Dupont LJ & Van Raemdonck DE (2006). Azithromycin Reduces Airway Neutrophilia and Interleukin-8 in Patients with Bronchiolitis Obliterans Syndrome. American Journal of Respiratory and Critical Care Medicine 174, 566–570.

24. Liu T, Zhang L, Joo D & Sun S-C (2017). NF-κB signaling in inflammation. Signal Transduction and Targeted Therapy 2, 17023.

25. Stellari FF, Sala A, Donofrio G, Ruscitti F, Caruso P, Topini TM, Francis KP, Li X, Carnini C, Civelli M & Villetti G (2014). Azithromycin inhibits nuclear factor-κB activation during lung inflammation: an in vivo imaging study. Pharmacol Res Perspect 2, e00058.

26. Haydar D, Cory TJ, Birket SE, Murphy BS, Pennypacker KR, Sinai AP & Feola DJ (2019). Azithromycin Polarizes Macrophages to an M2 Phenotype via Inhibition of the STAT1 and NF-κB Signaling Pathways. J Immunol 203, 1021–1030.

27. Murphy BS, Sundareshan V, Cory TJ, Hayes D, Jr, Anstead MI & Feola DJ (2008). Azithromycin alters macrophage phenotype. Journal of Antimicrobial Chemotherapy 61, 554–560.

28. Asbjarnarson A, Joelsson JP, Gardarsson FR, Sigurdsson S, Parnham MJ, Kricker JA & Gudjonsson T (2025). The Non-Antibacterial Effects of Azithromycin and Other Macrolides on the Bronchial Epithelial Barrier and Cellular Differentiation. Int J Mol Sci 26,

29. Halldorsson S, Gudjonsson T, Gottfredsson M, Singh PK, Gudmundsson GH & Baldursson O (2010). Azithromycin Maintains Airway Epithelial Integrity during Pseudomonas aeruginosa Infection. American Journal of Respiratory Cell and Molecular Biology 42, 62–68.

30. Kricker JA, Page CP, Gardarsson FR, Baldursson O, Gudjonsson T & Parnham MJ (2021). Nonantimicrobial Actions of Macrolides: Overview and Perspectives for Future Development. Pharmacological Reviews 73, 1404–1433.

31. Cuevas E, Huertas D, Montón C, Marin A, Carrera-Salinas A, Pomares X, García-Nuñez M, Martí S & Santos S (2023). Systemic and functional effects of continuous azithromycin treatment in patients with severe chronic obstructive pulmonary disease and frequent exacerbations. Front Med (Lausanne) 10, 1229463.

32. Southern KW & Barker PM Azithromycin for cystic fibrosis. European Respiratory Journal 24, 834–838.

33. Saiman L, Marshall BC, Mayer-Hamblett N, Burns JL, Quittner AL, Cibene DA, Coquillette S, Fieberg AY, Accurso FJ, Campbell III PW & Group ftMS (2003). Azithromycin in Patients With Cystic Fibrosis Chronically Infected With Pseudomonas aeruginosaA Randomized Controlled Trial. JAMA 290, 1749–1756.

34. Valery PC, Morris PS, Byrnes CA, Grimwood K, Torzillo PJ, Bauert PA, Masters IB, Diaz A, McCallum GB, Mobberley C, Tjhung I, Hare KM, Ware RS & Chang AB (2013). Long-term azithromycin for Indigenous children with non-cystic-fibrosis bronchiectasis or chronic suppurative lung disease (Bronchiectasis Intervention Study): a multicentre, double-blind, randomised controlled trial. The Lancet Respiratory Medicine 1, 610–620.

35. Hare KM, Grimwood K, Chang AB, Chatfield MD, Valery PC, Leach AJ, Smith-Vaughan HC, Morris PS, Byrnes CA, Torzillo PJ & Cheng AC (2015). Nasopharyngeal carriage and macrolide resistance in Indigenous children with bronchiectasis randomized to long-term azithromycin or placebo. European Journal of Clinical Microbiology & Infectious Diseases 34, 2275–2285.

36. Phaff SJ, Tiddens HAWM, Verbrugh HA & Ott A (2006). Macrolide resistance of Staphylococcus aureus and Haemophilus species associated with long-term azithromycin use in cystic fibrosis. Journal of Antimicrobial Chemotherapy 57, 741–746.

37. Southern KW, Solis-Moya A, Kurz D & Smith S (2024). Macrolide antibiotics (including azithromycin) for cystic fibrosis. Cochrane Database Syst Rev 2, Cd002203.

38. Taylor SL, Leong LEX, Mobegi FM, Choo JM, Wesselingh S, Yang IA, Upham JW, Reynolds PN, Hodge S, James AL, Jenkins C, Peters MJ, Baraket M, Marks GB, Gibson PG, Rogers GB & Simpson JL (2019). Long-Term Azithromycin Reduces Haemophilus influenzae and Increases Antibiotic Resistance in Severe Asthma. American Journal of Respiratory and Critical Care Medicine 200, 309–317.

39. Murray CJL, Ikuta KS, Sharara F, Swetschinski L, Robles Aguilar G, Gray A, Han C, Bisignano C, Rao P, Wool E, Johnson SC, Browne AJ, Chipeta MG, Fell F, Hackett S, Haines-Woodhouse G, Kashef Hamadani BH, Kumaran EAP, McManigal B, Achalapong S, Agarwal R, Akech S, Albertson S, Amuasi J, Andrews J, Aravkin A, Ashley E, Babin F-X, Bailey F, Baker S, Basnyat B, Bekker A, Bender R, Berkley JA, Bethou A, Bielicki J, Boonkasidecha S, Bukosia J, Carvalheiro C, Castañeda-Orjuela C, Chansamouth V, Chaurasia S, Chiurchiù S, Chowdhury F, Clotaire Donatien R, Cook AJ, Cooper B, Cressey TR, Criollo-Mora E, Cunningham M, Darboe S, Day NPJ, De Luca M, Dokova K, Dramowski A, Dunachie SJ, Duong Bich T, Eckmanns T, Eibach D, Emami A, Feasey N, Fisher-Pearson N, Forrest K, Garcia C, Garrett D, Gastmeier P, Giref AZ, Greer RC, Gupta V, Haller S, Haselbeck A, Hay SI, Holm M, Hopkins S, Hsia Y, Iregbu KC, Jacobs J, Jarovsky D, Javanmardi F, Jenney AWJ, Khorana M, Khusuwan S, Kissoon N, Kobeissi E, Kostyanev T, Krapp F, Krumkamp R, Kumar A, Kyu HH, Lim C, Lim K, Limmathurotsakul D, Loftus MJ, Lunn M, Ma J, Manoharan A, Marks F, May J, Mayxay M, Mturi N, Munera-Huertas T, Musicha P, Musila LA, Mussi-Pinhata MM, Naidu RN, Nakamura T, Nanavati R, Nangia S, Newton P, Ngoun C, Novotney A, Nwakanma D, Obiero CW, Ochoa TJ, Olivas-Martinez A, Olliaro P, Ooko E, Ortiz-Brizuela E, Ounchanum P, Pak GD, Paredes JL, Peleg AY, Perrone C, Phe T, Phommasone K, Plakkal N, Ponce-de-Leon A, Raad M, Ramdin T, Rattanavong S, Riddell A, Roberts T, Robotham JV, Roca A, Rosenthal VD, Rudd KE, Russell N, Sader HS, Saengchan W, Schnall J, Scott JAG, Seekaew S, Sharland M, Shivamallappa M, Sifuentes-Osornio J, Simpson AJ, Steenkeste N, Stewardson AJ, Stoeva T, Tasak N, Thaiprakong A, Thwaites G, Tigoi C, Turner C, Turner P, van Doorn HR, Velaphi S, Vongpradith A, Vongsouvath M, Vu H, Walsh T, Walson JL, Waner S, Wangrangsimakul T, Wannapinij P, Wozniak T, Young Sharma TEMW, Yu KC, Zheng P, Sartorius B, Lopez AD, Stergachis A, Moore C, Dolecek C & Naghavi M (2022). Global burden of bacterial antimicrobial resistance in 2019: a systematic analysis. The Lancet 399, 629–655.

40. Renna M, Schaffner C, Brown K, Shang S, Tamayo MH, Hegyi K, Grimsey NJ, Cusens D, Coulter S, Cooper J, Bowden AR, Newton SM, Kampmann B, Helm J, Jones A, Haworth CS, Basaraba RJ, DeGroote MA, Ordway DJ, Rubinsztein DC & Floto RA (2011). Azithromycin blocks autophagy and may predispose cystic fibrosis patients to mycobacterial infection. J Clin Invest 121, 3554–63.

41. Mukai S, Moriya S, Hiramoto M, Kazama H, Kokuba H, Che XF, Yokoyama T, Sakamoto S, Sugawara A, Sunazuka T, Ōmura S, Handa H, Itoi T & Miyazawa K (2016). Macrolides sensitize EGFR-TKI-induced non-apoptotic cell death via blocking autophagy flux in pancreatic cancer cell lines. Int J Oncol 48, 45–54.

42. Hodge S, Tran HB, Hamon R, Roscioli E, Hodge G, Jersmann H, Ween M, Reynolds PN, Yeung A, Treiberg J & Wilbert S (2017). Nonantibiotic macrolides restore airway macrophage phagocytic function with potential anti-inflammatory effects in chronic lung diseases. American Journal of Physiology-Lung Cellular and Molecular Physiology 312, L678–L687.

43. Glick D, Barth S & Macleod KF (2010). Autophagy: cellular and molecular mechanisms. J Pathol 221, 3–12.

44. Yu L, Chen Y & Tooze SA (2018). Autophagy pathway: Cellular and molecular mechanisms. Autophagy 14, 207–215.

45. Hu W, Chan H, Lu L, Wong KT, Wong SH, Li MX, Xiao ZG, Cho CH, Gin T, Chan MTV, Wu WKK & Zhang L (2020). Autophagy in intracellular bacterial infection. Seminars in Cell & Developmental Biology 101, 41–50.

46. Derendorf H (2020). Excessive lysosomal ion-trapping of hydroxychloroquine and azithromycin. Int J Antimicrob Agents 55, 106007.

47. Liao SX, Sun PP, Gu YH, Rao XM, Zhang LY & Ou-Yang Y (2019). Autophagy and pulmonary disease. Ther Adv Respir Dis 13, 1753466619890538.

48. Nakahira K, Pabon Porras MA & Choi AM (2016). Autophagy in Pulmonary Diseases. Am J Respir Crit Care Med 194, 1196–1207.

49. Hill C & Wang Y (2022). Autophagy in pulmonary fibrosis: friend or foe? Genes & Diseases 9, 1594–1607.

50. Zhang Y, Zhang J & Fu Z (2022). Role of autophagy in lung diseases and ageing. European Respiratory Review 31, 220134.

51. Mizumura K, Cloonan S, Choi ME, Hashimoto S, Nakahira K, Ryter SW & Choi AM (2016). Autophagy: Friend or Foe in Lung Disease? Ann Am Thorac Soc 13 **Suppl 1**, S40–7.

52. Lam HC, Cloonan SM, Bhashyam AR, Haspel JA, Singh A, Sathirapongsasuti JF, Cervo M, Yao H, Chung AL, Mizumura K, An CH, Shan B, Franks JM, Haley KJ, Owen CA, Tesfaigzi Y, Washko GR, Quackenbush J, Silverman EK, Rahman I, Kim HP, Mahmood A, Biswal SS, Ryter SW & Choi AM (2013). Histone deacetylase 6-mediated selective autophagy regulates COPD-associated cilia dysfunction. J Clin Invest 123, 5212–30.

53. Monick MM, Powers LS, Walters K, Lovan N, Zhang M, Gerke A, Hansdottir S & Hunninghake GW (2010). Identification of an autophagy defect in smokers’ alveolar macrophages. J Immunol 185, 5425–35.

54. Luciani A, Villella VR, Esposito S, Brunetti-Pierri N, Medina D, Settembre C, Gavina M, Pulze L, Giardino I, Pettoello-Mantovani M, D’Apolito M, Guido S, Masliah E, Spencer B, Quaratino S, Raia V, Ballabio A & Maiuri L (2010). Defective CFTR induces aggresome formation and lung inflammation in cystic fibrosis through ROS-mediated autophagy inhibition. Nature Cell Biology 12, 863–875.

55. Quarrington RD, Sapula SA, Lester SE, Miller MM, Kos VM, Kopp BT, Jersmann HP, Blencowe A & Roscioli E (2024). Modification of Azithromycin to Mitigate its Arrest of Autophagy. bioRxiv 2024.04.25.591217.

56. Janas A & Przybylski P (2019). 14- and 15-membered lactone macrolides and their analogues and hybrids: structure, molecular mechanism of action and biological activity. European Journal of Medicinal Chemistry 182, 111662.

57. ISO I (2019). 20776-1: 2019. Susceptibility Testing of Infectious Agents and Evaluation of Performance of Antimicrobial Susceptibility Test Devices—Part 1: Broth Micro-Dilution Reference Method for Testing the In Vitro Activity of Antimicrobial Agents Against Rapidly Growing Aerobic Bacteria Involved in Infectious Diseases. Broth micro-dilution reference method for testing the in vitro activity of antimicrobial agents against rapidly growing aerobic bacteria involved in infectious diseases

58. Kaja S, Payne AJ, Naumchuk Y & Koulen P (2017). Quantification of Lactate Dehydrogenase for Cell Viability Testing Using Cell Lines and Primary Cultured Astrocytes. Curr Protoc Toxicol 72, 2.26.1–2.26.10.

59. Farha MA, Tu MM & Brown ED (2025). Important challenges to finding new leads for new antibiotics. Current Opinion in Microbiology 83, 102562.

60. Poh W-P, Anthony K, E. LS, T. NP, O. BL, N. RP, Sandra H & and Roscioli E (2020). COPD-Related Modification to the Airway Epithelium Permits Intracellular Residence of Nontypeable Haemophilus influenzae and May Be Potentiated by Macrolide Arrest of Autophagy. International Journal of Chronic Obstructive Pulmonary Disease 15, 1253–1260.

61. Takano N, Hiramoto M, Yamada Y, Kokuba H, Tokuhisa M, Hino H & Miyazawa K (2023). Azithromycin, a potent autophagy inhibitor for cancer therapy, perturbs cytoskeletal protein dynamics. Br J Cancer 128, 1838–1849.

62. Zhang L, Song L, Liu Z, Li H, Lu Y, Li Z & Ma S (2010). Synthesis and antibacterial activity of novel 3-O-carbamoyl derivatives of clarithromycin and 11,12-cyclic carbonate azithromycin. European Journal of Medicinal Chemistry 45, 915–922.

63. Yan M, Ma X, Dong R, Li X, Zhao C, Guo Z, Shen Y, Liu F, Ma R & Ma S (2015). Synthesis and antibacterial activity of 4″-O-(trans-β-arylacrylamido)carbamoyl azithromycin analogs. European Journal of Medicinal Chemistry 103, 506–515.

64. Talman S, Uzun S, Djamin RS, Baart SJ, Grootenboers M, Aerts J & van der Eerden M (2021). Long-Term Azithromycin Maintenance Treatment in Patients with Frequent Exacerbations of Chronic Obstructive Pulmonary Disease. Int J Chron Obstruct Pulmon Dis 16, 495–498.

65. Kragol G, Steadman VA, Marušić Ištuk Z, Čikoš A, Bosnar M, Jelić D, Ergović G, Trzun M, Bošnjak B, Bokulić A, Padovan J, Glojnarić I & Eraković Haber V (2022). Unprecedented Epimerization of an Azithromycin Analogue: Synthesis, Structure and Biological Activity of 2’-Dehydroxy-5″-Epi-Azithromycin. Molecules 27,

66. Balloy V, Deveaux A, Lebeaux D, Tabary O, le Rouzic P, Ghigo JM, Busson PF, Boëlle PY, Guez JG, Hahn U, Clement A, Chignard M, Corvol H, Burnet M & Guillot L (2014). Azithromycin analogue CSY0073 attenuates lung inflammation induced by LPS challenge. Br J Pharmacol 171, 1783–94.

67. Hoppenbrouwers T, Bastiaan-Net S, Garssen J, Pellegrini N, Willemsen LEM & Wichers HJ (2022). Functional differences between primary monocyte-derived and THP-1 macrophages and their response to LCPUFAs. PharmaNutrition 22,

68. Chanput W, Mes JJ & Wichers HJ (2014). THP-1 cell line: an in vitro cell model for immune modulation approach. Int Immunopharmacol 23, 37–45.

69. Madhvi A, Mishra H, Leisching GR, Mahlobo PZ & Baker B (2019). Comparison of human monocyte derived macrophages and THP1-like macrophages as in vitro models for M. tuberculosis infection. Comp Immunol Microbiol Infect Dis 67, 101355.

70. Paul SM, Mytelka DS, Dunwiddie CT, Persinger CC, Munos BH, Lindborg SR & Schacht AL (2010). How to improve R&D productivity: the pharmaceutical industry’s grand challenge. Nature Reviews Drug Discovery 9, 203–214.

71. Rousseaux CG & Bracken WM (2013). Chapter 21 - Overview of Drug Development. In Haschek and Rousseaux’s Handbook of Toxicologic Pathology (Third Edition), edn, ed. Haschek WM, Rousseaux CG & Wallig MA, 647–685. Academic Press, Boston.

72. Tang B & More V (2023). Recent Advances in Drug Discovery Toxicology. International Journal of Toxicology 42, 535–550.

